## Supporting information for "Tracking and perceiving diverse motion signals: Directional biases in human smooth pursuit and perception"

### Supporting information: Further exploratory analyses between perceptual subgroups

The following materials provide details on the further exploratory analyses conducted: In the exploratory analyses, we showed that observers potentially have different perceptual bias patterns (Fig 8). In addition, perceptual biases correlated with pursuit biases (Fig 7). Because pursuit bias was dynamic and appeared to be stronger during the initial pursuit phase (Fig 5 a and b), we further explored whether observers with assimilation perceptual biases consistently showed a larger pursuit bias than observers with contrast perceptual biases over time. If observers with assimilation perceptual biases showed a larger pursuit bias consistently over time, it might reflect a general response tendency in these observers. By contrast, if the difference in pursuit bias between participants with different perceptual biases was developed during later pursuit phases, there might be a later perceptual modulation on pursuit. In addition, as the direct consequence of eye movements is the change in the retinal image, we also examined how net motion energy of retinal image changed over time, and whether it differed between the perceptual subgroups.

#### S1. Temporal dynamics of pursuit bias between perceptual subgroups

To explore the temporal dynamics of pursuit bias in observers with different perceptual bias patterns, we calculated the average pursuit direction bias in three time windows (S1 Fig a): the initiation phase (from pursuit onset to 140 ms after pursuit onset), the early steady-state phase (the first half of steady-state phase), and the late steady-state phase (the second half of steady-state). We conducted a two-way rmANOVA of pursuit bias with *subgroup* (assimilation and contrast) and *pursuit phase* (initiation, early steady-state, and late steady-state) as factors. We found that compared to the contrast group, the assimilation group had a slower decrease in pursuit bias over time (S1 Fig b), indicated by a significant *subgroup* × *pursuit phase* interaction effect [$F(2, 36) = 4.88$, $p = 0.01$, $\eta_p^2 = 0.21$]. Post-hoc *t*-tests with Tukey-adjusted *p* values showed that pursuit bias did not differ between the assimilation and contrast groups during the initiation [$t(54) = -1.43$, $p = 0.71$, 95% CI of difference = (-2.66, 0.93)] and early steady-state [$t(54) = 2.22$, $p = 0.24$, 95% CI of difference = (-0.45, 3.14)] phases. The assimilation group had a stronger pursuit bias than the contrast group only during the late steady-state phase [$t(54) = 3.15$, $p = 0.03$, 95% CI of difference = (0.12, 3.70)]. In fact, some observers from the contrast group even showed a negative pursuit bias during the late steady-state phase (S1 Fig b). Overall, the difference in pursuit biases between people having different perceptual bias patterns developed over time.

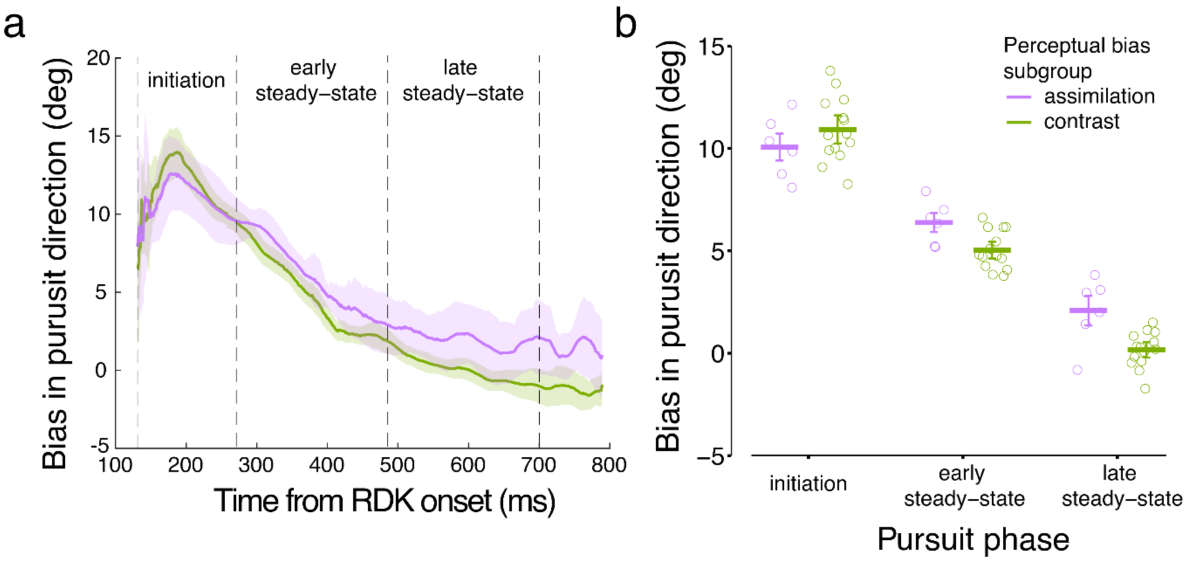


**S1 Fig.** Biases in perceptual subgroups over time. (a) Color indicates the perceptual bias subgroup, see legends in panel (b). Solid lines indicate the mean pursuit bias in each subgroup. Shaded areas indicate the 95% CI. Dashed vertical lines indicate time points of the pursuit onset, and the start, middle point, and end of the steady-state phase analysis window. (b) Biases in pursuit direction in perceptual subgroups across the three pursuit phases. Horizontal bars indicate the mean across observers. Error bars indicate the 95% CI. Circles indicate the mean of individual observers.

#### S2. Net motion energy between perceptual subgroups

During eye movements, both retinal and extraretinal signals are required to recover the actual object motion in the world. As people with different perceptual biases tended to have various magnitudes of pursuit biases, it would be interesting to see how retinal image motion was affected. In several studies using RDK stimuli, a small proportion of observers perceived motion in the opposite direction of the RDK, even when the RDK had 100% motion coherence [1]. The cause of such individual differences is unclear. One explanation is that even 100%-coherence RDKs could have motion energy [2] in the opposite direction [1,3]. In the current study, motion energy of the retinal image may not directly correspond to pursuit direction biases, since the retinal image motion is affected by both speed and direction of the eyes. Therefore, we calculated the motion energy of retinal images over time, to examine whether motion energy of the stimuli could be a source of the diverse motion perception. If our stimuli contain motion energy in the opposite direction of internal motion, a diversity in perception might be encouraged.

We calculated the averaged bias in the net motion energy of retinal images for each observer across time. Specifically, for each trial we first generated the 2D retinal image of the dots across time. Then, two pairs of spatiotemporal filters were convolved with the retinal image to calculate the net motion energy over time. We adopted the demo code from Mather [4] and used the spatiotemporal parameters detailed in a previous study [5]. Specifically, each pair of the spatiotemporal filter was the sum of different combinations of two spatial and two temporal filters. Each pair was selective for either the upward or the downward direction. The spatial filters were even and odd symmetric fourth-order Cauchy functions:

$$f_1(x, y) = cos^4(\alpha)\, cos(4\alpha)\, exp\left(-\frac{x^2}{2\sigma_g^2}\right)$$

$$f_2(x, y) = cos^4(\alpha)\, sin(4\alpha)\, exp\left(-\frac{x^2}{2\sigma_g^2}\right)$$

where $\alpha = tan^{-1}(y/\sigma_c)$, $\sigma_c = 0.35°$, and $\sigma_g = 0.05°$. The half width of the spatial filters was 0.7°. Two temporal filters were defined by the following functions:

$$g_1(t) = (60t)^3\, exp(-60t)\left[\frac{1}{3!} - \frac{(60t)^2}{(3+2)!}\right]$$

$$g_2(t) = (60t)^3\, exp(-60t)\left[\frac{1}{5!} - \frac{(60t)^2}{(5+2)!}\right]$$

The duration of the temporal filters was chosen to be nine frames, about 106 ms. Since the pursuit latency was 132 ± 13 ms in the current study, 106 ms was roughly enough time to gather information for pursuit planning/updating. It was also comparable to the temporal filter length used in a previous study for pursuit and perception [6]. $f_1g_1 + f_2g_2$ and $f_1g_2 - f_2g_1$ (element-wise multiplication of $f$ and $g$) would pass information in the upward direction, whereas $f_1g_2 + f_2g_1$ and $f_1g_1 - f_2g_2$ would pass information in the downward direction. After convolving each filter with the 3D spatiotemporal retinal image pattern, the results of each pair were squared and summed up to calculate the motion energy in the upward and downward direction separately. Then, motion energy in the downward direction was subtracted from that in the upward direction to yield the net motion energy.

The net motion energy was normalized between −1 and 1. A positive bias value indicates more motion energy in the internal motion direction. Surprisingly, we found that our stimuli contained slight motion energy in the opposite direction to internal motion, as shown by the negative bias values during the initial duration before pursuit onset (S2 Fig). With the eyes moving, more motion energy in the opposite direction was observed. The net motion energy of retinal images did not seem to differ between perceptual subgroups, although there seemed to be a trend that the net motion energy of retinal images for the assimilation group is less in the opposite direction to internal motion (S2 Fig).

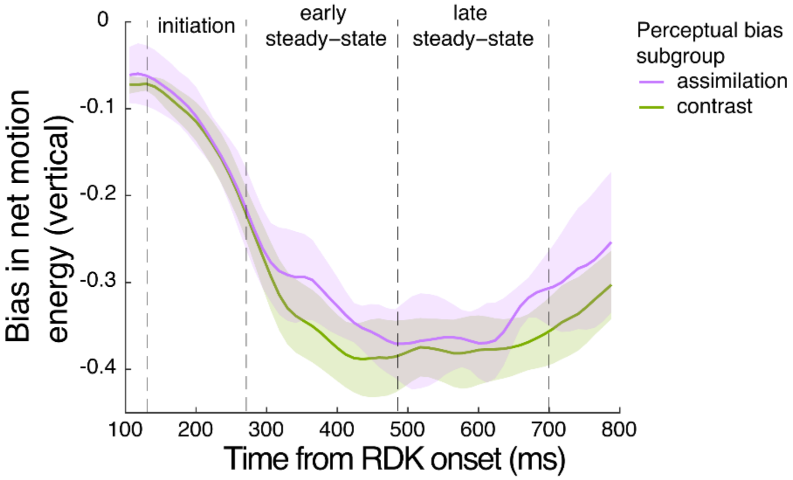


**S2 Fig. Biases in net motion energy in the vertical dimension between perceptual subgroups.** Positive values indicate that there was more motion energy in the same direction as internal motion. Solid lines indicate the mean of each perceptual subgroup. Shaded areas indicate the 95% CI. Dashed vertical lines indicate time points of the pursuit onset, and the start, middle point, and end of the steady-state phase analysis window.
